# Genomic insertion sequence populations of an ancient chemosynthetic symbiont

**DOI:** 10.64898/2026.11.25.769167

**Authors:** Olivia L. Mathieson, Nicolas C. H. Schröder, Anna Mankowski, Julian S. T. Kiefer, Nikolaus Leisch, Jessie L. Maier, Christer Erséus, Bruno Huettel, Jörg Ott, Florian Scharhauser, Nicole Dubilier, Harald R. Gruber-Vodicka, Manuel Kleiner

**Author notes:** Correspondence: Olivia L. Mathieson, Manuel Kleiner.

## Abstract

Genetic isolation as a consequence of host restriction of bacterial symbionts can lead to reductive genome evolution resulting in small, functionally streamlined genomes that are biased toward high AT contents. Frequently, reductive genome evolution leads to the loss of transposable elements, which are one of the major initial drivers of symbiont genome evolution through massive expansion events following initial host restriction. Under specific circumstances, where host-restricted bacteria continue to experience genetic mixing, transposable elements are retained in and, in some cases, introduced to symbiont genomes, even in ancient symbioses. Very little is known about these transposable element populations, regarding density, diversity, and activity as well as their influence on symbiont evolution and physiology. Here, we explored genomic population dynamics of insertion sequence (IS) elements, a type of transposable element, in 647 genomes of the chemosynthetic Gammaproteobacteria symbiont *Candidatus* Thiosymbion from three distantly related marine invertebrate hosts. We observed host-group specific patterns of genomic properties, likely reflective of symbiont genome evolution driven by varying degrees of genetic isolation due to differences in symbiont localization (ecto- to endosymbiont). We found that *Thiosymbion* genomes had highly diverse and dynamic IS populations, which varied by both symbiont localization and symbiont lineage with instances of clade-specific gain, loss, and expansion of certain IS families. Combined, our results suggest that variable IS expansion in *Thiosymbion* likely contributes to high intraspecific variability in genome structure.

## Introduction

Exploring drivers of bacterial genome evolution is critical for understanding how bacteria adapt to novel environments (e.g., a host). One of the major drivers of microbial genome evolution is mobile genetic elements (Siguier et al., 2014; Werren, 2011). In fact, it has been estimated that genes encoding transposases, the enzymes responsible for movement of many types of mobile genetic elements, are among the most abundant genes on the planet (Aziz et al., 2010). Insertion sequence (IS) elements are among the simplest and most ubiquitous mobile genetic elements, consisting of a transposase coding region flanked by inverted repeats (Siguier et al., 2015, 2014). Though simple in structure, IS elements are quite diverse and can exhibit unique ecological and evolutionary dynamics within host genomes (Siguier et al., 2015, 2014).

Like many biological entities IS elements are classified into families that vary in terms of transposase chemistry, characteristics of flanking inverted repeats, and carriage of additional coding regions, but also in regulation, insertion activity, genomic niche, and ultimately physiological consequences for their host genome (Guo et al., 2023; Kanai et al., 2023; Siguier et al., 2014). IS elements are often referred to as “selfish elements” (Werren et al., 1988), due to their occasionally deleterious effects. These effects are often considered the reason for their low abundance in most bacterial genomes and their tight regulation preventing frequent movements. Some IS families have been associated with specific regulatory and physiological functions, interactions with other mobile genetic elements, and environmental adaptation (e.g., Arashida et al., 2022; Cassier-Chauvat et al., 1997; Fordham et al., 2022; Razavi et al., 2020; Tempel et al., 2022; Vandecraen et al., 2017; Wedel et al., 2023). Despite their prevalence and growing list of potential roles, exploring genomic IS population dynamics from an ecological perspective is a relatively new approach to studying these mobile genetic element populations and is limited, especially in diverse systems (Gaudillière-Jami et al., 2026; Guo et al., 2023; Kanai et al., 2023; Siguier et al., 2014).

In closed genetic systems, such as genetically isolated endosymbionts, IS populations have been proposed to undergo a period of expansion and subsequent elimination from the genome without the genetic mixing necessary to replenish these populations (Plague et al., 2008). In such symbionts, we also expect certain patterns of genome evolution that follow host restriction, namely reduction in genome size and GC content which may be driven by bursts of transposable element expansion and deleterious homologous recombination (Moran and Plague, 2004; Plague et al., 2008). Characterizing these processes has been biased toward very small symbiont genomes of terrestrial insects, such as the genome of the pea aphid symbiont *Buchnera aphidicola* (616-642 kbp) and several other vertically transmitted, intracellular insect symbionts (Moran and Bennett, 2014), as well as chemosynthetic symbionts of marine animals such as *Candidatus* Ruthia magnifica (1.16 Mbp, 34% GC content; Roeselers et al., 2010), *Ca.* Vesicomyosocius okutanii from deep-sea clams (1.02 Mbp, 32% GC content; Kuwahara et al., 2008, 2007) and *Ca.* Riegeria from meiofauna flatworms (1.34 Mbp, 51.8% GC content; Jäckle et al., 2019). However, it has been shown that transmission mode (i.e., horizontal, vertical, or mixed) and exposure to genetic mixing during transmission may disrupt evolutionary processes attributed to bottle-neck effects and preserve genome size and mobile genetic element populations in the genome (Lund et al., 2014; Russell et al., 2020, 2017). For example, Lund et al. (2014) suggested that the vertically transmitted, extracellular earthworm symbiont *Verminephrobacter* retains high densities of IS elements in its genome over long evolutionary times (>60 million years) due to genetic mixing with co-occurring symbionts as well as with environmental bacteria during transmission (Bright and Bulgheresi, 2010). In systems such as *Verminephrobacter*, genome erosion and AT-bias seem to be avoided and genomic populations of IS elements are maintained alongside signs of bottle-neck driven genome evolution (e.g., higher substitution rates and low codon usage bias) (Lund et al., 2014).

*Candidatus* Thiosymbion (hereafter *Thiosymbion*), a globally distributed, chemosynthetic symbiont, has been associated with marine invertebrate hosts for more than 100 million years (Mankowski et al., 2026). *Thiosymbion* is associated with three marine worm host groups (gutless oligochaetes and nematodes) where it exhibits host-specific localizations that span the full range of symbiotic localizations (Zimmermann et al. 2016). Stilbonematinae-associated *Thiosymbion* are ectosymbionts which exhibit a high degree of host specificity but are exposed to the surrounding environment (Ott et al., 2004; Scharhauser et al., 2024, 2020), gutless oligochaete-associated *Thiosymbion* are subcuticular endosymbionts that frequently occur in a microbial consortium of up to six other endosymbionts (Dubilier et al., 2005, 2001; Mankowski et al., 2026; Woyke et al., 2006), and *Astomonema*-associated *Thiosymbion* are internal endosymbionts that are presumably genetically isolated as they occur either as intracellular endosymbionts (Musat et al., 2007; Ott et al., 1982) or as extracellular endosymbionts in the center of the worm potentially representing the lumen of the modified gut (Giere et al., 1995). Though the evolutionary distance between *Thiosymbion* hosts is vast, *Thiosymbion* do not generally form monophyletic clades based on these host groupings. While Stilbonematinae-associated *Thiosymbion* form host genus-specific clades, *Thiosymbion* of gutless oligochaetes do not form host group-specific clades and instead seem to have independently arose several times potentially from host-switches from Stilbonematinae (Zimmermann et al., 2016). In contrast, *Astomonema*-associated *Thiosymbion* are monophyletic based on current sampling efforts (Zimmermann et al., 2016). Interestingly, *Thiosymbion* of *Olavius algarvensis* (gutless oligochaete) has been shown to have high IS element densities in its genome and high transposase expression detected at the protein level (Kleiner et al., 2013; Woyke et al., 2006), suggesting potential transposase activity and possible IS movement on a micro-evolution scale. The density and diversity of IS populations in *Thiosymbion* from other hosts and how IS populations interact with symbiont evolution remain largely unresolved.

Here, we explore 647 *Thiosymbion* genomes from three globally distributed marine invertebrate host groups (Stilbonematinae, gutless oligochaetes, and *Astomonema*) to investigate the interactions between genome evolution and genomic IS element populations and how this interacts with genetic isolation on the spectrum from ectosymbiont to intracellular endosymbiont. Using these data we assessed the features of genomic evolution (e.g., genome size, GC content, and coding density) and how this varied by host group. We describe IS populations in symbiont genomes and correlate these populations with genomic properties to assess potential interactions between IS elements and genomic evolution. Finally, we construct a phylogenetic tree containing symbionts from all three host groups and three free-living relatives to explore phylogenetic patterns associated with genomic evolution and IS populations. Combined, we reveal how the host-associated lifestyle of a symbiont promotes specific dynamics between IS populations and genomic evolution across a gradient of genetic mixing, potentially providing insights into micro-evolutionary dynamics across diverse host associated lifestyles.

## Materials & Methods

### Description of genomes

We used 647 *Thiosymbion* metagenome assembled genomes (MAGs) sourced from 429 gutless oligochaetes, 182 Stilbonematinae, and 36 *Astomonema* individuals collected from global sampling sites between 1991 and 2026 (**Supplementary Data S1**). Samples were preserved in the field by freezing in a liquid nitrogen dry shipper or at -80℃ or fixed in 80% ethanol or RNAlater (Thermo Fisher Scientific, Waltham, MA, USA). We extracted DNA with the DNeasy Blood & Tissue Kit (Qiagen, Hilden, Germany). Some genomes were sequenced on Illumina platforms through the US Department of Energy Joint Genome Institute (Berkeley, CA, USA; previously published Mankowski et al., 2026). Additional genomes were sequenced through the Max Planck Genome Centre (Cologne, Germany) on Illumina platforms in 2 x 100, 150, or 250 bp paired-end read mode. DNA concentration was increased using the MinElute PCR Purification kit (Qiagen) when needed. Ultra-low input DNA libraries were prepared using the Ovation Ultralow Library System V2 kit (NuGEN or, later, Tecan Genomics) and low-input DNA libraries were prepared with the NEBNext Ultra™ DNA v2 Library Prep Kit for Illumina (New England Biolabs). When necessary for the pipelines used, initial DNA fragmentation to 400 bp fragment size was done with an S2 device from Covaris. Library preparation was performed following manufacturer’s instructions. If possible, with respect to input DNA size and amount, a miniature Tn5 transposase-based protocol (Nextera-LITE; Rowan et al., 2019) with a limited PCR step was applied. Sequencing method for each genome is available in **Supplementary Data S2**.

Genome assembly and binning of 168 *Thiosymbion* genomes from gutless oligochaetes (Mankowski et al., 2026) and 1 genome (Lonei1, NCBI: ASM435995v1) from Stilbonematinae were previously reported. We assembled all additional genomes using the same pipeline. In brief, reads were trimmed and error corrected using bbduk v39.08 (Bushnell, 2014) and the BayesHammer SPAdes v3.15.5 module (Bankevich et al., 2012; Nikolenko et al., 2013; Prjibelski et al., 2020) before assembly with MEGAHIT v1.2.9 (Li et al., 2015) and binning with MetaBAT v2.16 (Kang et al., 2015), MaxBin v2.2.6 (Wu et al., 2014), and CONCOCT v1.1.0 (Alneberg et al., 2014). The best bins from these were selected with DAStool v1.1.6 (Sieber et al., 2018). The resulting bins were checked for quality using CheckM v1.2.2 (Parks et al., 2015) and CheckM2 v1.0.1 (Chklovski et al., 2023) and taxonomically classified using GTDB-Tk v2.3.2 (Chaumeil et al., 2020). We also included the genomes of three free-living relatives of *Thiosymbion* – MASp6, ERR10921342_A, and SRR31168895_A. MASp6 is a previously published MAG sourced from salt marsh sediments colonized by *Spartina alterniflora* and is available under the NCBI BioSample identifier SAMN20958563 (Thomas et al., 2014). We used Sandpiper v2.0 (Woodcroft et al., 2026) to mine publicly available metagenomes for datasets containing *Candidatus* Thiosymbion or close relatives (GTDB genus g Thiosymbion). We processed the metagenomes as described above and binned a candidate MAG from two datasets: the Tara-Pacific coral samples from Palau (Planes et al., 2019; NCBI SRA: ERR10921342) and metagenomes of stony coral tissue loss disease from Dry Tortugas National Park, Florida, USA (NCBI BioProject Accession: PRJNA1177872, NCBI SRA: SRR31168895). We named the MAGs for the sequencing run identifier, ERR10921342_A and SRR31168895_A, respectively. We calculated genome statistics (size, GC content, coding density) using Bakta (v. 1.12.1 with full v. 6.0 database; Schwengers et al., 2021) and seqkit (Shen et al., 2024) (**Supplementary Data S3**).

### Symbiont phylogeny

To generate the symbiont phylogeny, we dereplicated genomes using dRep (v3.7.1; Olm et al., 2017) at the species level to remove redundant genomes leading to 151 dereplicated genomes and 3 outgroup genomes. To construct a phylogeny from shared single-copy genes, we used GToTree (v1.6.34; Lee, 2019) with 159 shared genes from the 172 genes in the Gammproteobacteria reference set to produce alignments and IQTree (v2.0.3; Nguyen et al., 2015) using the model finder feature to select the optimal model. Branch supports were calculated using ultrafast bootstrapping and SH-like approximate ratio test, both with 1000 replicates. The tree was visualized using iTOL (v7.6; Letunic and Bork, 2024) using free-living relatives as the outgroup to root the tree.

### IS element Identification

To identify and quantify IS elements, we used ISEScan (v1.7.3; Xie and Tang, 2017). Due to high fragmentation in some of our genome assemblies, likely due to high density of IS elements (Woyke et al., 2006), we retained both complete and partial IS elements identified by this tool. Because the genome assemblies were highly fragmented, we further classified partial insertion sequences as “edge-cases” if they were located within 250 bp of a contig edge.

### Analyses of genome properties, insertion sequence diversity and density

IS density was calculated by taking percentage of base pairs associated with IS elements relative to the total number of base pairs in the symbiont genome. Diversity of the IS population was characterized using IS family and cluster classifications (herein referred to interchangeably as “IS subgroups” or “IS clusters”) output by ISEScan. Statistical analyses were performed in R (v4.2.2). Diversity metrics (i.e., richness, evenness, Morisita-Horn dissimilarity) were calculated at the family and cluster level of the IS population. To identify possible relationships between genomic properties and IS density, richness, and evenness, we used Spearman’s rank correlation (sm_statCorr function in smplot2 v0.2.6) (**Supplementary Table 1**). We used a Kruskal-Wallis test to explore the relationship between host group and other variables (length, GC content, density, IS density, richness, and evenness; **Supplementary Table 2**), followed by a Dunn post-hoc test with Benjamini-Hochberg FDR correction for pairwise comparison. We explored the independent impact of host group, symbiont species, sampling region, and genomic properties (length, GC content, coding density, and IS density) on IS population composition using adonis2 (vegan v2.6-4; Oksanen et al., 2026). We compared inter- and intraspecies dissimilarity regarding IS element diversity by comparing pairwise dissimilarity distances within and between symbiont species. To assess correlation between the IS element diversity dissimilarity matrix and *Thiosymbion* phylogeny, we performed Mantel tests on a Spearman correlation with 10,000 permutations (vegan v2.6-4; Oksanen et al., 2026).

## Results

### Symbionts follow host group-specific genomic evolution trajectories

We analyzed 647 genomes of the Gammaproteobacteria symbiont genus *Thiosymbion*, encompassing a total of 151 symbiont species, and 3 free-living relatives. The completeness of the genomes ranged from 75-100% with contamination below 10%. We found a large amount of variability in genome characteristics between *Thiosymbion* genomes. Sizes of *Thiosymbion* genomes ranged from 2.17 to 7.80 Mbp. GC content also had a broad range from 43 to 63% as well as coding density which ranged from 48 to 83%. In comparison, the genome sizes of the free-living relatives ranged from 3.64 to 4.57 Mbp with an average GC content of 61.27% (± 0.31% SD) and an average coding density of 82.30% (± 1.31% SD). Unexpectedly, we observed a moderate negative correlation between genome size and GC content (r = -0.44, p < 0.001; Spearman’s rank correlation; **Fig. 1A**), i.e. lower GC content in larger genomes, and a weak positive correlation between genome size and coding density (r = 0.18, p < 0.001; Spearman’s rank correlation; **Fig. 1B**). We found that symbiont host group (*Astomonema*, Stilbonematinae, and gutless oligochaetes) interacted with the relationship between genome size and GC content or coding density (**Fig. 1A-B**). Generally, we observed that the genomes from Stilbonematinae ectosymbionts and gutless oligochaete endosymbionts followed similar trends, with the genomes from *Astomonema* endosymbionts following inverse trends or exhibiting negligible trends. For example, we observed a moderate negative correlation between genome size and GC content in the Stilbonematinae ectosymbiont and gutless oligochaete endosymbiont genomes (r = -0.66, p < 0.001 and r = -0.60, p < 0.001, respectively; Spearman’s rank correlation), while we observed a moderate positive correlation between genome size and GC content in genomes of *Astomonema* endosymbionts (r = 0.41, p = 0.014; Spearman’s rank correlation). Genome size, GC content, and coding density significantly differed between symbiont host group, with gutless oligochaete and *Astomonema* endosymbionts having smaller genomes on average compared to Stilbonematinae ectosymbionts and free-living relatives (**Fig. 1C**). *Astomonema* endosymbionts had both the lowest GC content and coding density (**Fig. 1D-E**).

**Figure 1.**
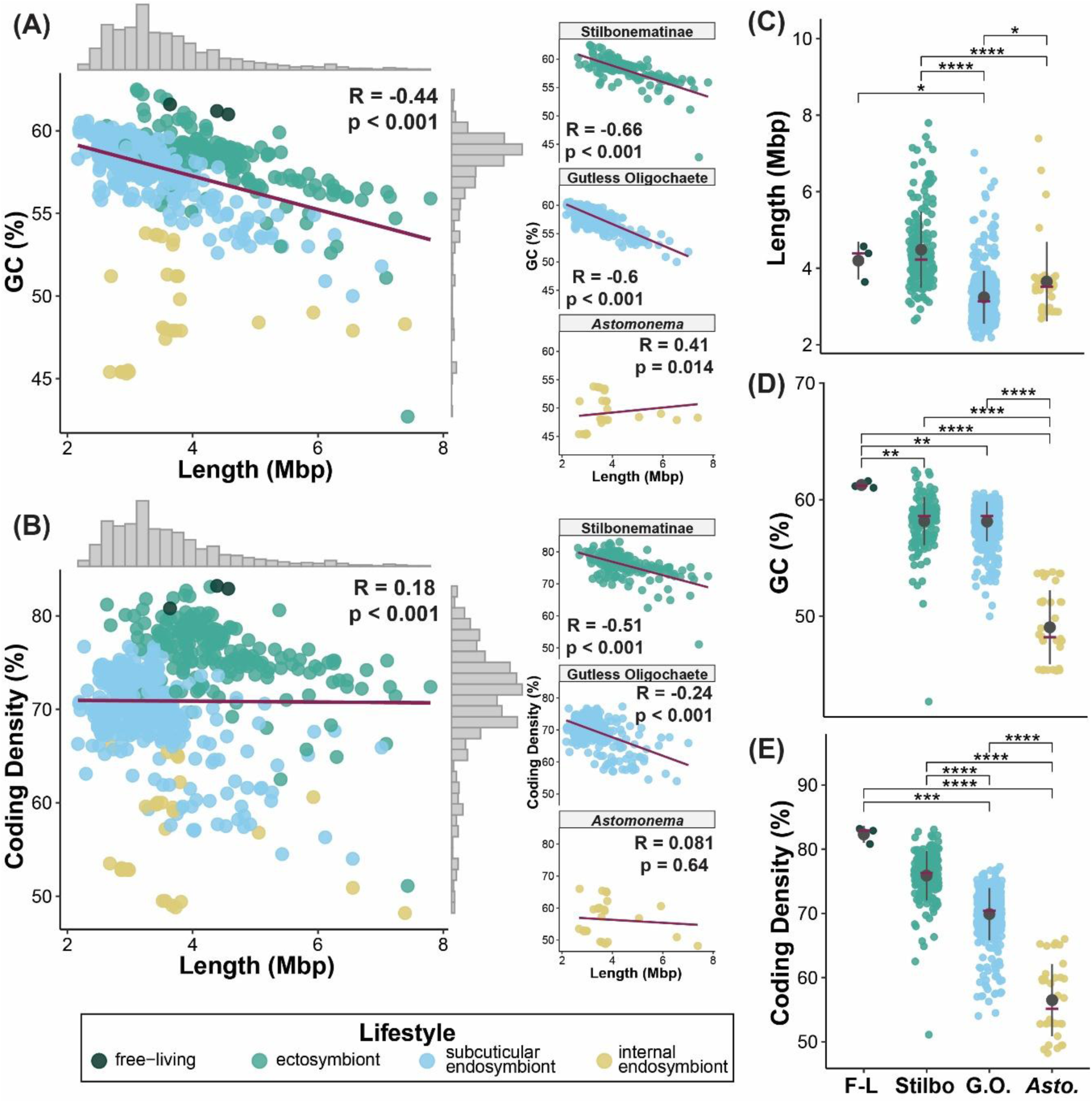
Interactions between *Thiosymbion* lifestyle and genomic properties. Interaction between genome length (Mbp) and (A) GC content (%) and (B) coding density (%) in 647 of *Thiosymbion* genomes and 3 free-living relatives (left) and interactions within host group subsets: *Astomonema* (n = 36), Stilbonematinae (n = 182), and gutless oligochaetes (n = 429) (right). Results of a Spearman’s rank correlation are printed on the graph (R = Spearman’s rho) and linear regression is represented by the red line. Data distribution is indicated by the histograms (grey bars) in the margins of the graphs. (C-E) Genome assembly size, GC content, and coding density across host groups. Mean is indicated by grey dot with error bars representing standard deviation and median is indicated by the red cross bar. Statistical significance is indicated by asterisks and cross bars and was determined by a Kruskal-Wallis test followed by Dunn’s post hoc test with Benjamini Hochberg correction (****: p.adj < 1e-04, ***: p.adj < 0.001, **: p.adj <0.01, *: p.adj < 0.05).

### *Thiosymbion* genomes can harbor large and diverse IS populations

We identified 67,240 IS elements from 24 IS families, as well as putative IS elements not yet assigned to IS families and from uncharacterized families across all 650 genomes (*Thiosymbion* and free-living relatives). This included 7,708 complete and 59,532 partial IS elements, as classified by ISEScan. We classified 34,131 of the partial IS elements as “edge cases” based on their proximity to edges of contigs (**Supplementary Fig. S1A**). Edge case and partial IS elements made up the largest component of identified IS elements in each genomic population with respectively 53 (range: 0-602) and 39 (range: 3-745) sequences per genome on average, while complete IS elements were much lower at 12 sequences per genome on average (range: 0-821) (**Supplementary Fig. 1B**). Due to genomic features that might interfere with adequate complete versus partial classification by ISEScan, such as high levels of fragmentation in some genome assemblies due to the high number of IS elements coupled with potentially complex IS architecture, we included complete, partial, and edge case IS elements in our quantification and diversity analyses. Combined this dataset is likely representative of the IS diversity in the global gutless oligochaete- and Stilbonematinae-associated *Thiosymbion* populations, as indicated by plateauing of the IS subgroup accumulation curve, but additional genomes from *Astomonema*-associated *Thiosymbion* would be necessary to capture the true diversity IS populations in this lineage (**Fig. 2A**). IS elements from the families IS5, IS630, IS66, IS1, IS481, and IS4 were the most abundant (**Fig. 2C**). In contrast, other IS elements were more sparse, often present in single occurrences in a few genomes.

**Figure 2.**
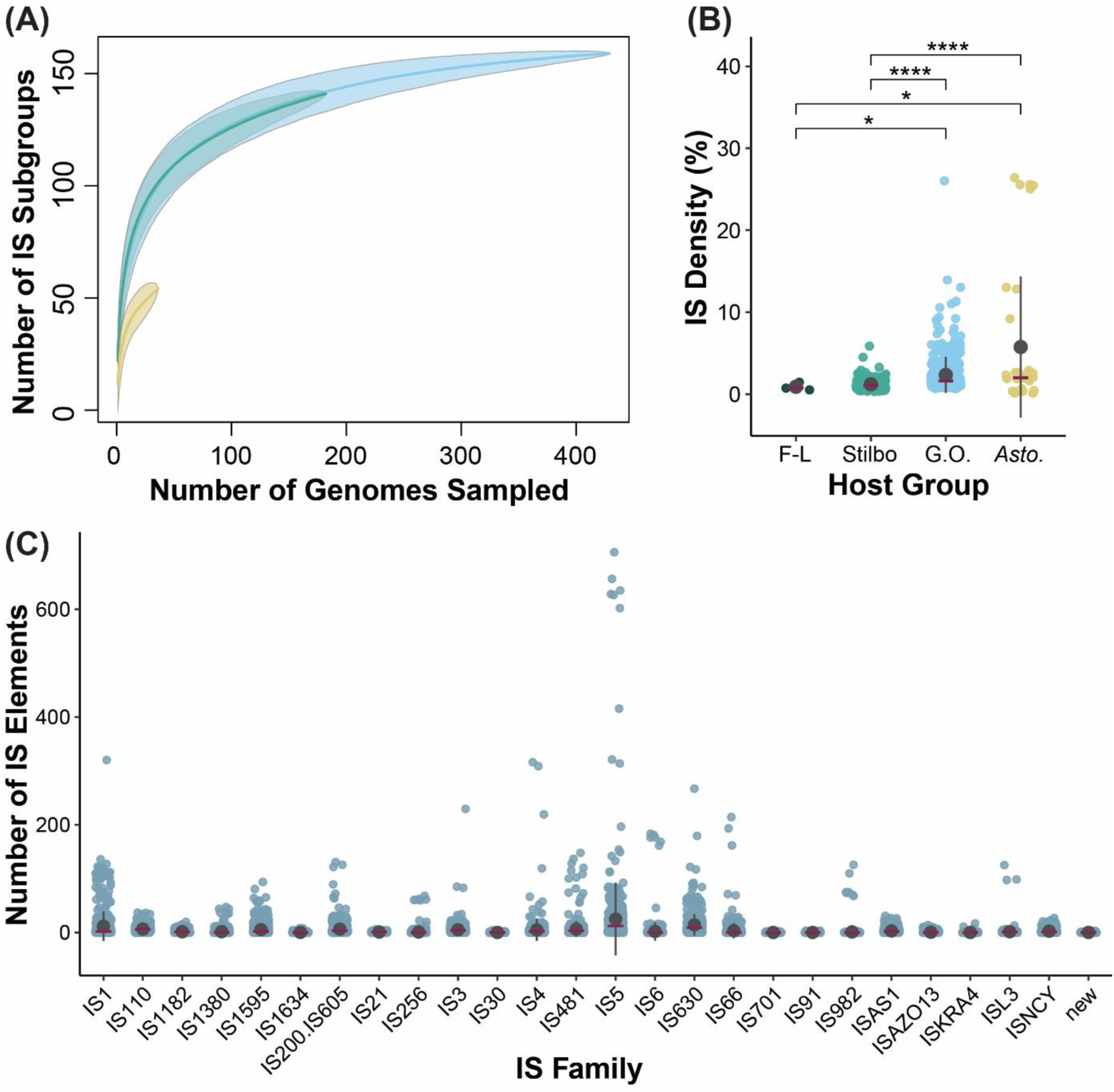
Insertion sequence element (IS) content of *Thiosymbion* genomes. (A) IS subgroup (cluster, as defined by ISEScan) accumulation curve. The line represents the mean accumulation curve from random permutations (permutations = 1000) of the data. Yellow represents *Astomonema* symbionts, blue represents gutless oligochaete symbionts, and green represents Stilbonematinae symbionts. (B) IS density per *Thiosymbion* genome across host groups given as % of the total number of nucleotides in the genome. The mean is indicated by a grey dot with error bars representing standard deviations and the median is indicated by the red cross bar. Statistical significance is indicated by asterisks and cross bars and was determined by a Kruskal-Wallis test followed by Dunn’s post hoc test with Benjamini-Hochberg correction (****: p.adj < 1e-04, ***: p.adj < 0.001, **: p.adj <0.01, *: p.adj < 0.05). (C) Number of IS elements detected per IS family and genome as assigned by ISEScan, again with mean indicated by a grey dot with error bars representing standard deviations and the median indicated by the red cross bar.

On average, genomes of symbionts had higher densities of IS elements than genomes of the free-living relatives. *Thiosymbion* of gutless oligochaetes and *Astomonema* had the highest IS loads (**Fig. 2B**). Free-living relatives, ectosymbionts (Stilbonematinae), and subcuticular endosymbionts (gutless oligochaetes) had higher richness (number of IS subgroups) and evenness (similarity of abundance distribution across IS subgroups) of the IS population (**Fig. 3A-B**). To understand how much IS populations vary between *Thiosymbion* genomes, we calculated a weighted (considering both IS subgroup and its abundance) dissimilarity matrix (Morisita-Horn Distances). We observed that within species IS population dissimilarity (average: 0.18, range: 9.9E-05 - 0.95, SD: 0.11) is lower than between species IS population dissimilarity (average: 0.73, range: 8.78E-03 - 1.0, SD = 0.18), suggesting that IS populations within species are more similar than between species, with gutless oligochaete and *Astomonema* endosymbionts exhibiting lower intraspecies IS population dissimilarity than Stilbonematinae ectosymbionts (**Fig. 3C**). Within species IS population dissimilarity varied from species to species (**Supplementary Fig. S2B-D**); however, the differences observed in intraspecies diversity between species did not seem to be driven by differences in number of genomes per species in our dataset. For example, a symbiont species for which we have 160 genomes has average dissimilarity of 0.19 while the highest intraspecies dissimilarity (0.64) is found in a symbiont species for which we only have 6 genomes (**Supplementary Fig. S2**). IS population composition, as measured by dissimilarity, significantly varied at both the IS family- and IS subgroup-level based on symbiont host group-associated lifestyle, symbiont species, and geographic sampling region (p < 0.001, PERMANOVA; **Fig. 3D, Supplementary Table 3**). We also found that genome length, GC content, coding density, and IS density significantly associated with the observed variance in composition, though to a lesser extent than lifestyle, species, and geography (**Supplementary Table 3**).

**Figure 3.**
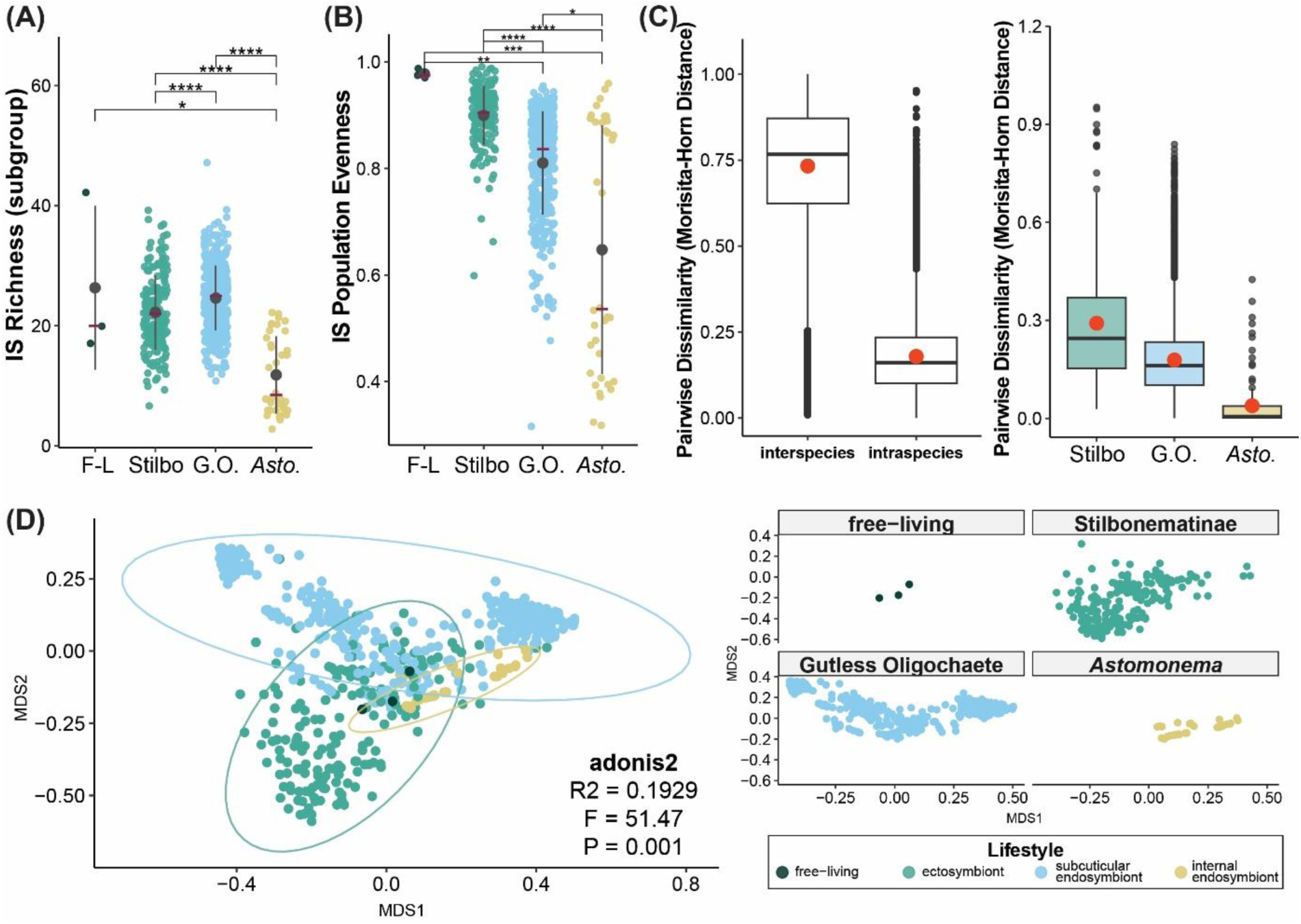
Insertion sequence population structure. (A) IS richness, or number of unique IS subgroups per genome, and (B) IS evenness at the subgroup-level across host group (Stilbo = Stilbonematinae; G.O. = gutless oligochaetes; Asto = *Astomonema*; F-L = free-living). Mean is indicated by a grey dot with error bars representing standard deviation and the median is indicated by the red cross bar. Statistical significance is indicated by asterisks and cross bars and was determined by a Kruskal-Wallis test followed by Dunn’s post hoc test with Benjamini-Hochberg correction (****: p.adj < 1e-04, ***: p.adj < 0.001, **: p.adj <0.01, *: p.adj < 0.05). (C) Inter- and intraspecies pairwise dissimilarity (left) and intraspecies pairwise dissimilarity within each host group (right). Each point is a pairwise distance between two IS populations from the same symbiont species (intraspecies) or different species (interspecies). In the boxplot, the center line indicates the group median and red dot indicates the group mean. The lower and upper portion of the box represents the first and third quartiles, while the whiskers extend to the minimum or maximum of the group or to 1.5x the interquartile range with data points falling outside of that represented as individual points. (D) MDS ordination of IS population (subgroup-level) based on Morisita-Horn Distances (left = full dataset; right = faceted by host group). Point color and ellipses (95% CI) indicate host-associated lifestyle.

### Interactions between IS populations and genomic properties differ between *Thiosymbion* from different host groups/lifestyles

We explored the relationship between genomic structural properties and IS population dynamics to understand how IS populations may be influencing *Thiosymbion* genome evolution. Overall, we observed only weak correlations between IS population metrics (density, richness, and evenness) and genomic properties (length, GC content, and coding density) when considering all 647 *Thiosymbion* and free-living relative genomes; however, these relationships varied by host group and, in some cases, inverted and stronger correlations were observed for specific host groups as compared to the correlations in the complete genome set (**Fig. 4**). For example, we observed a negative correlation between genome size, GC content, and coding density and IS density (**Fig. 4A-C**); however, when we looked within host group we observed stronger, positive correlations associated with *Thiosymbion* of *Astomonema*. This was most notable in the strong relationship between IS density and coding density in *Thiosymbion* from *Astomonema* (R = 0.89, p < 0.001; **Fig. 4C**). We observed similar distinction between *Thiosymbion* of *Astomonema* and other hosts when correlating IS richness and evenness, with the strongest trends observed in *Astomonema*-associated *Thiosymbion*. For example, GC content and coding density exhibited strong (R = 0.91, p < 0.001; **Fig. 4E**) and moderate (R = 0.6, p < 0.001; **Fig. 4F**) positive correlations with IS richness in *Astomonema*-associated *Thiosymbion*, respectively. Overall, larger genomes appear to support IS populations with greater richness and evenness, but this trend also varied between host groups (**Fig. 4D,G**). For example, *Thiosymbion* of gutless oligochaetes had the strongest relationship between genome size and richness (R = 0.5, p < 0.001), while *Thiosymbion* of *Astomonema* had the strongest relationship between genome size and evenness (R = 0.61, p < 0.001) (**Fig. 4D,G**). Interestingly, the relationships between stilbonematine *Thiosymbion* IS population structure and genomic structural properties are overall weaker than those observed in the other two host groups.

**Figure 4.**
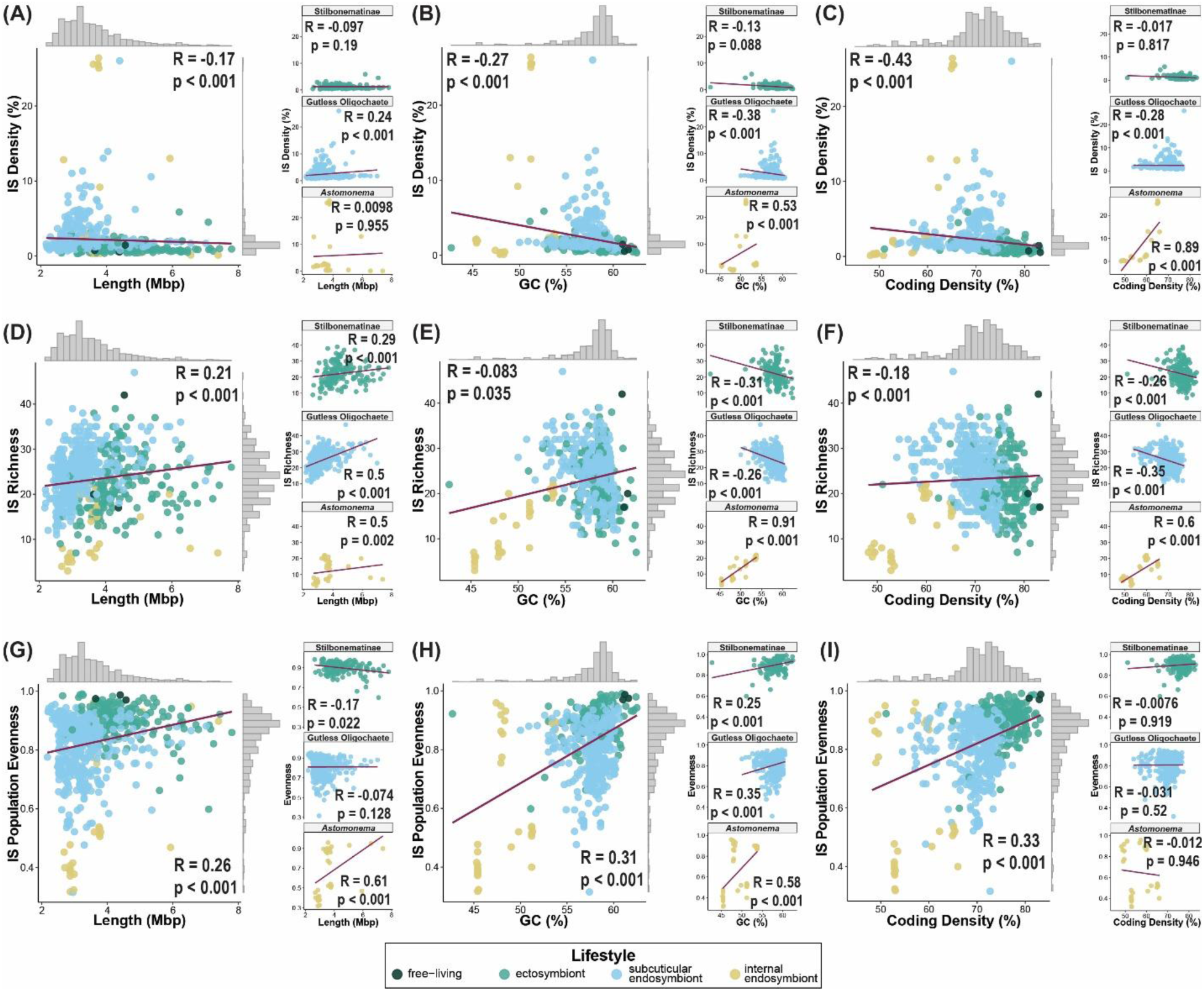
Correlation between IS population and genomic properties. (A-C) IS density (%), (D-F) IS richness and (G-I) IS evenness at the subgroup-level correlated with genome assembly length (Mbp), GC content, and coding density. Results of Spearman’s rank correlations are printed on the graph. R (Spearman’s rho) indicates strength and direction of the correlation and p indicates the significance of the correlation. A linear regression trendline is represented by the red line. Each panel contains the full population of *Thiosymbion* genomes included in this study (left) and correlation within host group subsets (right). Distribution of each variable is shown in the histograms (grey bars) in the margins of the graphs.

### Intraspecies IS population dissimilarity suggests IS element movement on micro-evolutionary scale

We constructed a phylogenomic tree with representative genomes from each of the 151 symbiont species and 3 free-living relatives using single copy genes of Gammaproteobacteria (**Fig. 5**). Most branches were supported at greater than 95% based on 1000 bootstraps. The tree revealed three large clades of *Thiosymbion*: two distinctly dominated by Stilbonematinae symbionts with multiple gutless oligochaete symbiont emergence events and one containing symbionts from all three host groups but primarily gutless oligochaete and all *Astomonema* symbionts. Consistent with prior work by Zimmermann et al. (2016), *Thiosymbion* of Stilbonematinae seem to make up the most basal lineages. In several cases, gutless oligochaete-associated symbionts diverged from Stilbonematinae symbionts creating lineages with mixed host groups that suggest the emergence of gutless oligochaete-associated *Thiosymbion* from Stilbonematinae-associated *Thiosymbion* has occurred several times (**Fig. 5**). Also consistent with Zimmermann et al. (2016), the presumably genetically isolated *Astomonema* endosymbionts form a single clade only seeming to have emerged once based this set of genomes, though there are fewer genomes representing this host group relative to gutless oligochaete and Stilbonematinae symbionts (**Fig. 5**). The clade containing the *Astomonema* symbiont clade is unique as it appears to contain potentially four re-emergence events of Stilbonematinae from gutless oligochaete symbionts, these are the only occurrences of apparent Stilbonematinae symbiont re-emergence in this tree (**Fig. 5**).

**Figure 5.**
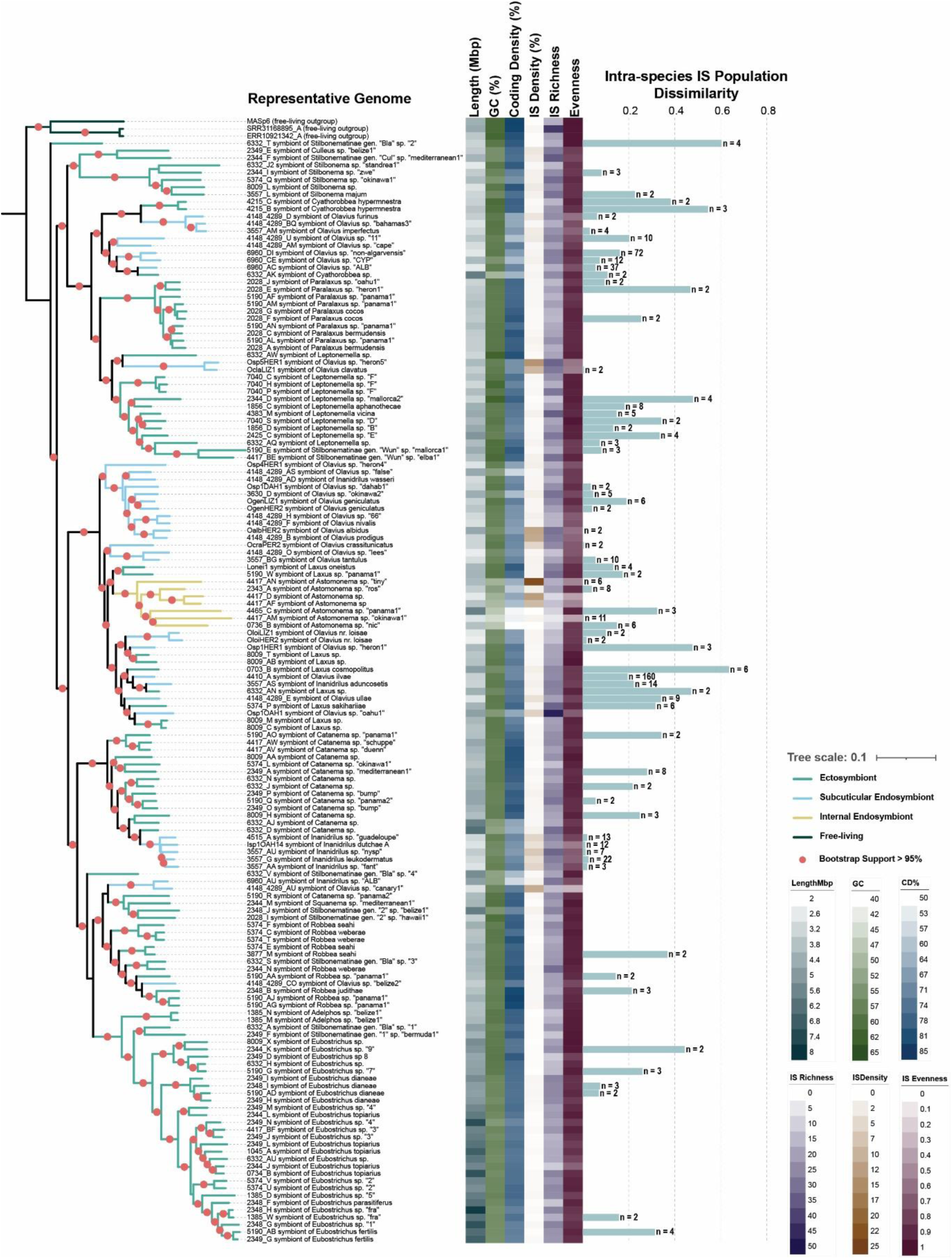
Phylogenomic tree of *Thiosymbion* showing genomic properties and IS population structure. The tree was constructed using a single representative genome for each species with the number of genomes available for computation of genome properties and intra-species comparisons indicated by ‘n’. Those without ‘n’ indicated are species with single genomes. Supports for nodes are indicated by a red dot (>95%; 1000 bootstraps). Branch colors represent host group. Average genome length, GC content, coding density, insertion sequence density, richness and evenness of subgroup-level genomic IS populations are indicated by gradient bars. Horizontal bars indicate average intraspecies IS population dissimilarity based on pairwise Morisita-Horn distances. Pairwise dissimilarity could not be calculated for species with single genome representatives.

Genome size, GC content, and coding density are distributed unevenly across the tree, with limited clade-level patterning (**Fig. 5**). Interestingly, one large clade of Stilbonematinae (genus *Eubostrichus*) *Thiosymbion* generally features genomes on that are consistently on the larger end of the spectrum, while other Stilbonematinae clades show more variation in genome size suggesting some phylogenetic signal in genome properties of Stilbonematinae *Thiosymbion*. Genome sizes of *Astomonema Thiosymbion* were highly variable while GC content and coding density were generally on the lower end. IS density was around 1% for most *Thiosymbion*, however, there were several subclades with clearly elevated IS densities. For example, the five members of the clade enclosed by *Thiosymbion* from *Inanidrilus* sp. “guadeloupe” and *Inanidrilus* sp. “fant” had IS densities around 5% (**Supplementary Fig. S3**). Generally, genomes with the highest IS densities belonged to *Thiosymbion* from gutless oligochaetes and *Astomonema Thiosymbion* and the clades in which the highest IS element densities were observed were all derived clades rather than ancestral clades. In some cases, the increased IS density appears to be driven by 1-2 dominating IS families and thus is frequently accompanied by lower IS population evenness and occasionally lower IS community richness (**Fig. 5, Supplementary Fig. S3**). For example, one *Astomonema*-associated *Thiosymbion* species exhibit remarkably high IS density (23.5% ± 5.16% SD), which appears to be driven by IS5 and IS6 family IS elements with 656 and 181 sequences in the representative genome for the species, respectively (**Fig. 2, Supplement Fig. S3**).

While symbiont species was a significant explanatory variable for the IS population dissimilarity in genomic IS population at the IS family- and subgroup-level (**Supplementary Table 3**), we did not detect association between the dissimilarity distances of IS populations of symbiont species representative genomes and the *Thiosymbion* phylogenomic tree at the IS family- or subgroup-level based on comparison between the phylogenomic distance matrix and IS population dissimilarity matrix (**Supplementary Fig S4-S5, Supplementary Table S4**; p > 0.9, Mantel). We did observe clustering of most *Astomonema*-associated *Thiosymbion* IS populations at the subgroup level and to some degree at the family level. This was not particularly surprising as *Astomonema*-associated *Thiosymbion* seemed to have fewer IS families in their IS populations compared to the remainder of the tree (**Supplementary Fig. S3**). Unique to *Astomonema*-associated *Thiosymbion* were two IS families, IS6 and IS982 (**Supplementary Fig. S3**). *Astomonema*-associated *Thiosymbion* genomes contained up to 5 subgroups of IS6, sharing only one of these subgroups with one of the free-living relatives. In contrast, only one subgroup of IS982 was identified and this family was entirely unique to *Astomonema-*associated *Thiosymbion*. These IS families were not consistently present across every *Astomonema-* associated *Thiosymbion* genome; however, when they were found they were typically in high abundance. *Astomonema*-associated *Thiosymbion*, were also notably missing prominent IS families that were present in the majority of *Thiosymbion*, e.g., IS1 and IS110 (**Supplementary Fig. S3**). Overall, there was variable patterning of IS populations across the full dataset, including within symbiont species. As mentioned above, within species dissimilarity at the subgroup-level varied from species to species and seems to be randomly distributed across the tree (**Fig. 5**). Notably, instances of high within species dissimilarity do not seem to follow a pattern related to IS density, richness, or evenness, and seem to be distributed independent of genome size, GC content, and coding density **(Fig. 5**).

## Discussion

Since symbiont host-associated lifestyle has been shown to influence the evolutionary trajectory of symbiont genomes and IS elements have been implicated as drivers of genomic evolution in symbionts broadly, we sought to explore the interaction between symbiont lifestyle (ectosymbionts, subcuticular endosymbionts, and internal endosymbionts), genome evolution, and IS population dynamics in a globally distributed symbiont genus across three host groups.

### Genome properties of *Thiosymbion* do not follow expectations for increasingly sexually isolated symbionts

It is broadly accepted that host restriction is a major evolutionary transition for bacteria, which is associated with specific, major changes in genome properties (McCutcheon and Moran, 2012; Moran and Plague, 2004). This is most intensely exemplified by symbionts that are sexually isolated because of their intracellular lifestyle and strict vertical transmission. These symbionts undergo predictable changes in their genome that result in streamlining of encoded functions, AT-bias, and overall reduction in genome size (McCutcheon and Moran, 2012; Moran and Bennett, 2014). Not every symbiont follows this trajectory, namely symbionts which undergo mixed or horizontal transmission, are exposed to the environment during vertical transmission, or symbionts exposed to other microbes in their host-associated lifestyle (i.e., associated with a consortium or microbiome) (Lund et al., 2014; Russell et al., 2020). In *Thiosymbion* we observed relatively large genomes for symbionts, with all genomes in this dataset being larger than 2 Mbp and with GC contents above 42% as compared to known sexually isolated symbionts of marine animals (e.g. chemosynthetic symbionts of deep-sea clams *Cand.* Ruthia magnifica and *Cand.* Vesicomyosocius okutanii, chemosynthetic symbionts of marine flatworms *Cand.* Riegeria) and terrestrial insects (e.g., *Buchnera aphidicola*, *Blochmannia floridanus*) with genome sizes ranging from 0.6 to 1.2 Mbp and GC content ranging from 25 to 52% (Gil et al., 2002; Jäckle et al., 2019; Kuwahara et al., 2008, 2007; McCutcheon and Moran, 2012; Moran and Bennett, 2014; Roeselers et al., 2010). This observation was in line with our expectations as most *Thiosymbion* symbionts are not sexually isolated, experiencing varied levels of exposure to the environment.

Interestingly, we found that genome properties varied by host group and did not always follow expected patterns. *Thiosymbion* of Stilbonematinae and gutless oligochaete followed similar trends for interactions between genome properties with symbionts from both host groups exhibiting negative correlation between genome length and both GC content and coding density, with *Thiosymbion* genomes from gutless oligochaete being on average smaller than that of *Thiosymbion* of Stilbonematinae. Since the *Thiosymbion* of gutless oligochaetes are subcuticular endosymbionts and more shielded from the environment than stilbonematine ectosymbionts (Dubilier et al., 2001; Mankowski et al., 2026; Ott et al., 2004; Woyke et al., 2006), this could fit within a model where genome reduction occurs upon increasing sexual isolation of the symbiont. In contrast, while *Thiosymbion* associated with gutless oligochaetes and Stilbonematinae still have access to external genetic material, in *Astomonema* we would expect complete or almost complete sexual isolation of the symbiont due to its intracellular or internally localized endosymbiotic lifestyle (Giere et al., 1995; Ott et al., 1982). Genomes of *Thiosymbion* in *Astomonema* were unexpectedly larger on average than the ones from gutless oligochaetes and showed a surprisingly low coding density of less than 60% on average (**Fig. 1C & E**), meaning the expected trend of genome reduction and associated increase of coding density was not observed for *Thiosymbion* of *Astomonema.* GC content was much lower in *Thiosymbion* of *Astomonema* as compared to the other two host groups, nearing 50% on average (**Fig. 1D**), but was higher than the very low GC contents described for sexually isolated symbionts of insects (20-44%; McCutcheon and Moran, 2012) and deep-sea clams (32-34%; Kuwahara et al., 2008, 2007; Roeselers et al., 2010). The GC content of *Astomonema*-associated *Thiosymbion* was similar to that of *Ca.* Riegeria santandreae, a vertically transmitted, intracellular endosymbiont of a marine flatworm with a reduced genome (1.3 Mbp) and GC content of 51.8%, suggesting that GC content in some symbionts experiencing genome reduction can remain relatively high (Jäckle et al., 2019). Indeed, one of the smallest symbiont genomes (0.14 Mbp) belonging to the insect alpha-proteobacteria symbiont *Hodgkinia cicadicola* has a remarkably high GC content of 58.4% (McCutcheon et al., 2009).

The misalignment between sexual isolation and expectations for resulting genome properties of *Astomonema*-associated *Thiosymbion* could potentially be explained in two ways. First, it is possible that *Thiosymbion* only recently (a few million years) associated with *Astomonema* and is currently undergoing the massive genome changes driven by sexual isolation that could ultimately result in gene loss, genome reduction, and potentially further reduction of GC content. The low coding density could be explained in this scenario by initial high activity and expansion of mobile elements, which is consistent with the IS population data (**Fig. 5**). The evolutionary age of the *Astomonema* symbiosis is currently unknown, and its determination would be critical to further establish support for this hypothesis. Second, potentially the transition of *Thiosymbion* within *Astomonema* between intracellular and extracellular endosymbiont lifestyles has triggered high rates of changes in genome structure, potentially supported by the variable IS density and diversity within the genomes of these symbionts. This could potentially explain the high variance in genome length with *Astomonema*-associated *Thiosymbion*.

Combined, our observations strongly support that host associated lifestyle is a critical determinant of the evolutionary trajectory of symbiont genomes and that this is likely linked to the genetic flux imposed by these lifestyle conditions on symbiont populations.

### IS element expansion in multiple derived *Thiosymbion* clades with some genomes reaching IS densities greater than 25%

One feature that may be contributing to the unique evolutionary dynamics of *Thiosymbion* may be the large and diverse IS populations that seem to occur in *Thiosymbion* genomes. It has been shown that symbiont genome evolution and reduction is promoted, in part, by mobilization and expansion of mobile genetic elements, which without exposure to genetic mixing ultimately lead to their own removal from the genome typically marking the end of large genetic changes in these systems (Moran and Plague, 2004), though this is not the case for every symbiont (Lund et al., 2014). In some cases, we observed that the IS populations of *Thiosymbion* were large with the most densely populated genomes containing over 1,000 IS elements, but this varied with IS density ranging from 0.1 to 26%. This is in line with previous reports that transposases make up more than 20% of genes encoded by *Thiosymbion* and that there are around 1,600 transposons in the genome of *Verminephrobacter*, a symbiont of a terrestrial oligochaete (Arumugaperumal et al., 2020; Woyke et al., 2006). In contrast, the lower IS density *Thiosymbion* genomes fall in line with the density of IS elements reported in other chemosynthetic Gammaproteobacteria symbionts of other marine invertebrates (deep sea clams, mussels, and annelids) which range from approximately 0-4% of the genome with symbionts transmitted by horizontal and mixed transmission having a higher percentage compared to those that were vertically transmitted (Coste et al., 2026).

In most cases, even in genomes with low IS density, the populations of IS elements were diverse, including a range of 2 to 20 IS elements families per genome. While we found that IS populations in *Thiosymbion* genomes were variable, the composition of IS populations of *Thiosymbion* seemed to be most similar within symbiont species, host group, and sampling location (**Fig. 3, Supplementary Table S3**). This was not surprising and it is likely that the effects of these variables are connected, as *Thiosymbion* species are associated with only one host group (Stilbonematinae, gutless oligochaete, or *Astomonema*) and mostly a single host species within a single sampling location.

Instances of high IS density appear to be driven by dominance of specific IS families in *Thiosymbion*. We observe this very clearly in *Astomonema* endosymbiont with high IS density where dominance of 2 or 3 IS families is typically coupled with decrease in overall IS population evenness. This might be in line with expectations for the beginning of a genome streamlining scenario, where selection relaxes allowing for expansion of certain IS groups leading to production of pseudogenes and ultimately genome reduction (Moran and Plague, 2004; Plague et al., 2008). In support of this, we observe a moderate positive correlation between IS population evenness and genome length, suggesting high evenness of the IS population is correlated with larger, more stable genomes. High IS density is not only associated with *Astomonema*-associated *Thiosymbion*, we also observe high IS density in some genomes of *Thiosymbion* in gutless oligochaete. These are typically associated with clades that have their most recent bifurcation with a Stilbonematinae symbiont. In these cases, it is unclear what is contributing to the ‘bursts’ of specific IS families in these IS populations. IS transposition is promoted by many stresses from starvation and changes in substrate availability to environmental triggers, like temperature stress and metal toxicity (Hall, 1999; Lee et al., 2020; Ohtsubo et al., 2005; Shin et al., 2023; Twiss et al., 2005; Vandecraen et al., 2017). Perhaps, these seemingly random bursts in the gutless oligochaete *Thiosymbion* are associated with an environmental trigger that was captured by the vast geographic and temporal ranges of this study.

### Variability of within symbiont species IS populations suggests high transpositional activity of IS elements on microevolutionary scales

Not every symbiont species had multiple genome representatives, but for those that did, we observed varying degrees of dissimilarity between genomic IS populations suggesting transpositional activity of IS elements within *Thiosymbion* populations on microevolutionary scales. We did not observe any clade-specific patterns associated with IS population dissimilarity, and events of high intraspecies IS element population dissimilarity occurred across the symbiont tree and within symbionts from all three host groups (**Fig. 5**) suggesting that the broad host grouping may not influence IS element activity. Variance in IS population dissimilarity across symbiont species could indicate that transpositional activity levels differ between *Thiosymbion* species. We speculate instances of high intraspecific IS population dissimilarity could reflect recent invasions or losses of IS elements, changes in environment impacting regulation of transposases, or other genomic disturbances that might lead to changes in selection pressures on specific genes allowing for IS expansion. Ultimately, this transpositional activity may lead to highly variable genome structures between *Thiosymbion* strains from separate individuals of the same species. Large intraspecific variation in IS populations has been previously reported in vertically transmitted symbionts of weevils, where it was reported that an IS element can have greater than 10-fold intraspecific copy number variation with one element (IS256) estimated to range 1,000 to 10,000 copies in field collected samples (Dougherty and Plague, 2008). While *Thiosymbion* does not have instances of 10,000 IS elements, we observe variability across our dataset on a similar scale at the IS family level (e.g., IS3 family varies from 3-85 sequences in *Thiosymbion* of *Laxus cosmopolitus*, IS256 family varies from 0-20 sequences in *Thiosymbion* of *Olavius* sp. “heron1”). While IS population variation is not a direct measure of IS mobility, it strongly suggests that IS transposition is occurring on microevolutionary scales in some *Thiosymbion* species.

### Limitations

In this study, we focused on characterizing the IS populations of *Thiosymbion* genomes and, therefore, chose to be inclusive of potential IS identifications over strict conservative estimates. In other words, we considered both complete and partial IS elements predicted by ISEscan to encompass complete and pseudogenized IS elements, as well as IS elements disrupted due to short read assembly methods. IS element population sizes are potentially over- or underestimated, because short-read based assembly method struggle to assemble highly repetitive genetic elements. Further, the classifications of IS elements in this study relies on the assignments and assumptions of the ISEScan workflow (Xie and Tang, 2017), and further work will be necessary to curate identified IS elements within the established taxonomies of TnCentral (Ross et al., 2021) and ISFinder (Siguier et al., 2006). For example, our current framework differentiates IS elements to the cluster level (as defined by ISEScan) within IS element families, but does not differentiate potentially different IS elements within these clusters and thus cannot completely accurately trace if an IS element represents an instance of an element that has been vertically inherited from a *Thiosymbion* ancestor or if it is a newly acquired IS element within a family/cluster already represented with other elements in *Thiosymbion* genomes.

## Conclusion

We observed host group-specific patterns of symbiont genome size, GC content, and coding density, likely reflective of host group-specific lifestyles of the symbiont. In many cases, symbiont genomes carried large and diverse IS populations in their genomes. Interactions of IS population dynamics with genomic properties appeared to be much lower in Stilbonematinae and gutless oligochaete symbionts as compared to *Astomonema* symbionts, potentially related to lack or decreased genetic isolation in these systems. In contrast, we observed strong correlations between IS population diversity metrics and several genomic properties in *Astomonema* symbionts which are presumably genetically isolated endosymbionts. *Astomonema* symbionts exhibit distinctive patterns of potential loss and gain of certain IS families, reaching high densities of specific IS families, some of which are unique to *Astomonema* and potentially reflect horizontal acquisition of these IS families. Combined, our results suggest that IS element population dynamics in *Thiosymbion* are likely influenced by the level of genetic exposure of the symbiont, rather than strictly by host restriction, and further, that high intraspecific variability of IS populations suggests movement of elements on microevolutionary scales.

## Supplementary Data

Supporting Information: File contains supplementary figures and tables.

Supplementary Data: (S1) Overview of the MAGs used in this study and the GTDB-Tk taxonomic assignment. (S2) Sequencing methods for the MAGs used in this study. (S3) Properties of *Thiosymbion* genomes.

## Data Availability

Many of the MAGs used in this study were previously published (see Mankowski et al. 2026). Additional MAGs used in this study will be made publicly available prior to publication and will be provided on request.

## Author Contributions

Conceived the study - O.L.M., H.R.G.-V, N.D., M.K.; Data Curation - O.L.M., A.M., N.C.H.S., N.L., C.E., H.R.G.-V, M.K.; Methodology - O.L.M., N.C.H.S., A.M., J.S.T.K., N.L., J.L.M., C.E., B.H., J.O., F.S., R.R-M., H.R.G.-V, M.K.; Interpretation of results - O.L.M., N.D., H.R.G.-V, M.K.; Visualization - O.L.M.; Writing - original draft - O.L.M. & M.K.; Writing - review & editing - all authors; Supervision - N.D., H.R.G.-V, M.K.; Funding Acquisition - N.D., H.R.G.- V, M.K.

## Acknowledgments

We are thankful for sample collections and field assistance by A. Gruhl, A. Ansebo, A. Blazejak, A.-C. Kreutzmann, C. Lott, C. Bergin, C. Wentrup, D. M. Michellod, E. M. Sogin, E. Mejlon, E. Mueller, F. Warnecke, F. Wells, J. Wippler, J. Zimmermann, K. Worsaae, K. Halanych, B. K. B. Seah, L. Matamoros, L. Gustavsson, M. P. Schimak, M. Hadfield, M. Sadowski, M. Weber, O. Giere, O. Jäckle, N. Bekkouche, O. Gros, P. Reid, P. Bouchet, P. De Wit, R. Rosello-Mora, S. Wetzel, A. Niclas, S. Bulgheresi, S. Sommer, T. Enders, and Y. Sato. In addition, we thank the crews of the Meteor research cruise M92 and the Sonne research cruise SO147, as well as the staff at the Carrie Bow Cay Laboratory, the Heron Island Research Station, the HYDRA Institute Elba, the Mediterranean Institute for Advanced Studies, the Lee Stocking Island Research Station, the Little Darby Island Research Station, the Lizard Island Research Station, and the Okinawa Institute of Science and Technology for supporting our sampling campaigns. This is contribution number XXXX from the Smithsonian’s Caribbean Coral Reef Ecosystems Program. We also thank the Max Planck Genome Centre and the EMBL Heidelberg IT Services and HPC resources (https://doi.org/10.5281/zenodo.12785829).

