## Supporting Information for "Genomic insertion sequence populations of an ancient chemosynthetic symbiont"

### Supplementary Tables

**Table S1.** Results of Spearman's rank correlation between assembly length (Len_Mbp), GC content (GC), coding density, IS density, cluster-level IS richness (richness_c), and cluster-level IS evenness (evenness_c) in full dataset and host group (Stilbo = Stilbonematinae; GO = gutless oligochaetes; Asto = Astomonema).

| Dataset | X | Y | R | p |
|---|---|---|---|---|
| **Full** | **Len_Mbp** | **GC** | **-0.44** | **<0.001** |
| Stilbo | Len_Mbp | GC | -0.66 | <0.001 |
| GO | Len_Mbp | GC | -0.60 | <0.001 |
| Asto | Len_Mbp | GC | 0.41 | 0.014 |
| **Full** | **Len_Mbp** | **CodingDensity** | **0.18** | **<0.001** |
| Stilbo | Len_Mbp | CodingDensity | -0.51 | <0.001 |
| GO | Len_Mbp | CodingDensity | -0.24 | <0.001 |
| Asto | Len_Mbp | CodingDensity | 0.081 | 0.64 |
| **Full** | **Len_Mbp** | **ISDensity** | **-0.17** | **<0.001** |
| Stilbo | Len_Mbp | ISDensity | -0.097 | 0.19 |
| GO | Len_Mbp | ISDensity | 0.24 | <0.001 |
| Asto | Len_Mbp | ISDensity | 0.0098 | 0.955 |
| **Full** | **GC** | **ISDensity** | **-0.27** | **<0.001** |
| Stilbo | GC | ISDensity | -0.13 | 0.088 |
| GO | GC | ISDensity | -0.38 | <0.001 |
| Asto | GC | ISDensity | 0.53 | <0.001 |
| **Full** | **CodingDensity** | **ISDensity** | **-0.43** | **<0.001** |
| Stilbo | CodingDensity | ISDensity | -0.017 | 0.817 |
| GO | CodingDensity | ISDensity | -0.28 | <0.001 |
| Asto | CodingDensity | ISDensity | 0.89 | <0.001 |
| **Full** | **Len_Mbp** | **richness_c** | **0.21** | **<0.001** |
| Stilbo | Len_Mbp | richness_c | 0.29 | <0.001 |
| GO | Len_Mbp | richness_c | 0.5 | <0.001 |
| Asto | Len_Mbp | richness_c | 0.5 | 0.002 |
| **Full** | **GC** | **richness_c** | **-0.083** | **0.035** |
| Stilbo | GC | richness_c | -0.31 | <0.001 |

| | | | | |
|---|---|---|---|---|
| GO | GC | richness_c | -0.26 | <0.001 |
| Asto | GC | richness_c | 0.91 | <0.001 |
| **Full** | **CodingDensity** | **richness_c** | **-0.18** | **<0.001** |
| Stilbo | CodingDensity | richness_c | -0.26 | <0.001 |
| GO | CodingDensity | richness_c | -0.35 | <0.001 |
| Asto | CodingDensity | richness_c | 0.6 | <0.001 |
| **Full** | **Len_Mbp** | evenness_c | **0.26** | **<0.001** |
| Stilbo | Len_Mbp | evenness_c | -0.17 | 0.022 |
| GO | Len_Mbp | evenness_c | -0.074 | 0.128 |
| Asto | Len_Mbp | evenness_c | 0.61 | <0.001 |
| **Full** | **GC** | evenness_c | **0.31** | **<0.001** |
| Stilbo | GC | evenness_c | 0.25 | <0.001 |
| GO | GC | evenness_c | 0.35 | <0.001 |
| Asto | GC | evenness_c | 0.58 | <0.001 |
| **Full** | **CodingDensity** | **evenness_c** | **0.33** | **<0.001** |
| Stilbo | CodingDensity | evenness_c | -0.0076 | 0.919 |
| GO | CodingDensity | evenness_c | -0.031 | 0.52 |
| Asto | CodingDensity | evenness_c | -0.012 | 0.946 |

**Table S2.** Results of Kruskal-Wallis test on assembly length (Len_Mbp), GC content (GC), coding density, IS density, cluster-level IS richness (richness_c), and cluster-level IS evenness (evenness_c) across symbiont lifestyles.

| X | Y | statistic | df | p |
|---|---|---|---|---|
| Lifestyle | Len_Mbp | 238.45 | 3 | 2.06E-51 |
| Lifestyle | GC | 106.45 | 3 | 6.38E-23 |
| Lifestyle | CodingDensity | 315.99 | 3 | 3.44E-68 |
| Lifestyle | ISDensity | 122.64 | 3 | 2.09E-26 |
| Lifestyle | richness_c | 90.63 | 3 | 1.60E-19 |
| Lifestyle | evenness_c | 163.00 | 3 | 4.14E-35 |

**Table S3.** Results of PERMANOVA (adonis2) analysis on the influence of symbiont lifestyle, geographic region, symbiont species, assembly size (Len_Mbp), GC content, coding density (CD), and insertion sequence density (ISD) on genomic insertion sequence population composition at the cluster-level and family-level.

| **Cluster-level** | | | |
|---|---|---|---|
| **Variable** | **R2** | **F** | **P** |
| Lifestyle | 0.1929 | 51.47 | 0.001 |
| Region | 0.3150 | 8.316 | 0.001 |
| SymbiontSpecies | 0.9305 | 43.75 | 0.001 |
| Len_Mbp | 0.0658 | 45.66 | 0.001 |
| GC (%) | 0.0508 | 34.65 | 0.001 |
| CD (%) | 0.0399 | 26.91 | 0.001 |
| ISD (%) | 0.0340 | 22.78 | 0.001 |
| **Family-level** | | | |
| **Variable** | **R2** | **F** | **P** |
| Lifestyle | 0.2580 | 74.87 | 0.001 |
| Region | 0.3559 | 10.00 | 0.001 |
| SymbiontSpecies | 0.9629 | 84.77 | 0.001 |
| Len_Mbp | 0.1080 | 78.44 | 0.001 |
| GC (%) | 0.0721 | 50.32 | 0.001 |
| CD (%) | 0.0624 | 43.12 | 0.001 |
| ISD (%) | 0.0300 | 20.06 | 0.001 |

**Table S4.** Mantel statistic based on Spearman's rank correlation rho.

| IS dissimilarity distance | Permutations | Mantel statistic (r) | p-value |
|---|---|---|---|
| Family-level | 10,000 | -0.08009 | 0.995 |
| Cluster-level | 10,000 | -0.05318 | 0.933 |

### Supplementary Figures

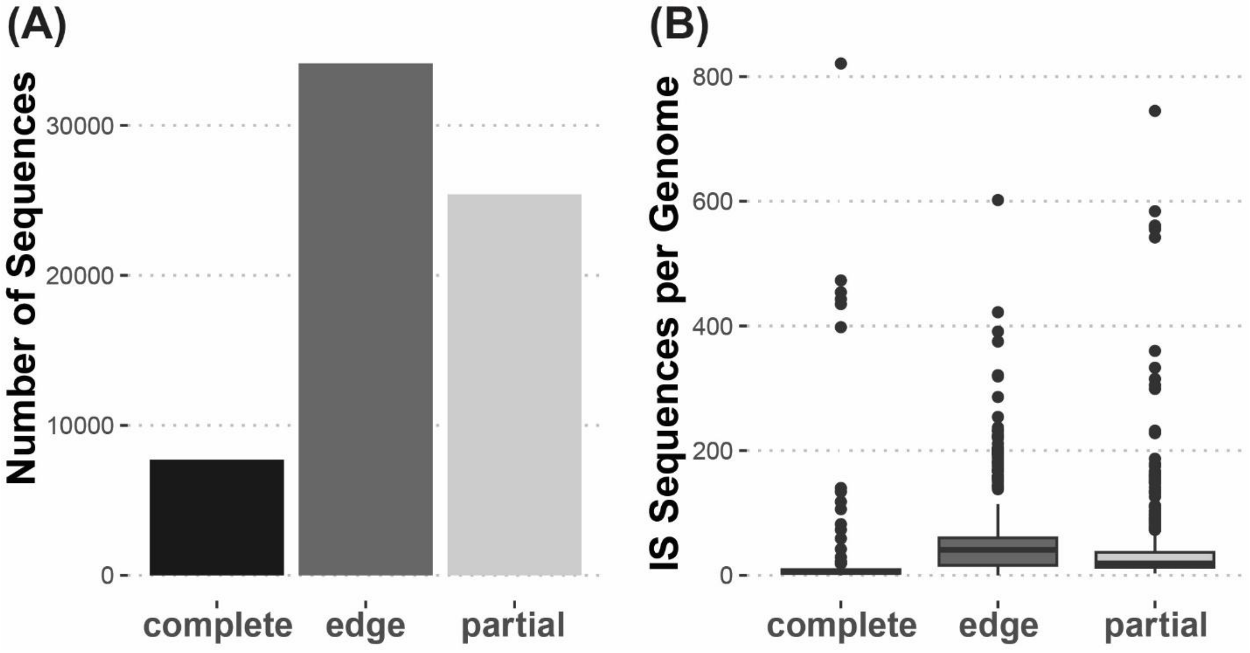


**Figure S1. Number of complete, edge-case, and partial IS elements identified in *Thiosymbion* genomes.** (A)The total number of complete, edge-case, and partial IS elements in 647 *Thiosymbion* and 3 free-living relative genomes. (B) Number of complete, edge-case, and partial IS elements per genome. The center line indicates the group median. The lower and upper portion of the box represents the first and third quartiles, while the whiskers extend to the minimum or maximum of the group or to 1.5x the interquartile range with data points falling outside of that represented as individual points.

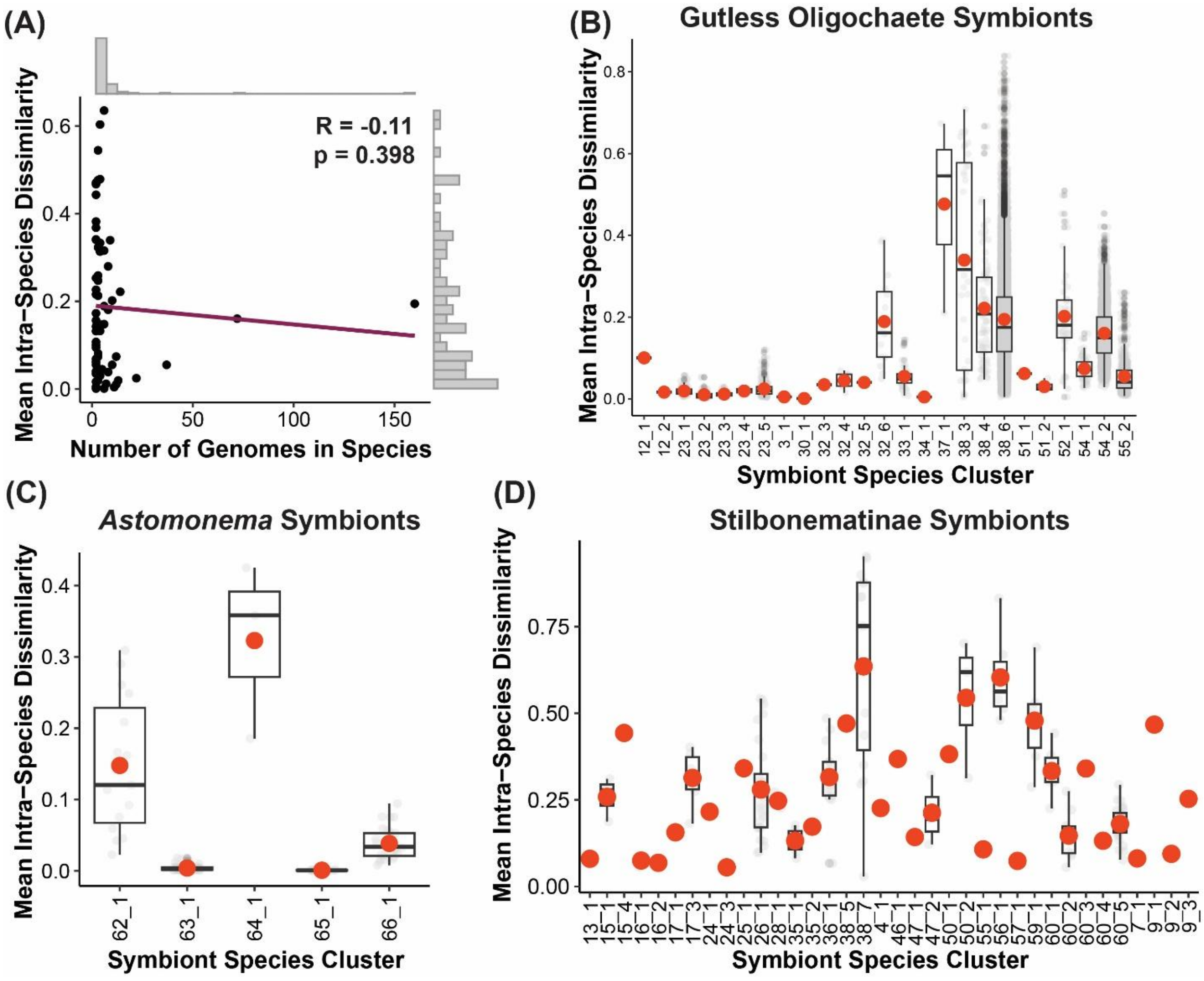


**Figure S2. Intraspecies IS population dissimilarity.** (A) Correlation between number of genomes per species and intraspecies dissimilarity (Morisita-Horn Distance). Results of a Spearman's rank correlation are printed on the graph (R = Spearman's rho) and the resulting trend is represented by the red line. (B-D) Intraspecies pairwise dissimilarity of gutless oligochaete (B), Stilbonematinae (C), and *Astomonema* (D) *Thiosymbion*. Each point is a pairwise distance between two IS populations from the same symbiont species. In the boxplot, the center line indicates the group median and red dot indicates the group mean. The lower and upper portion of the box represents the first and third quartiles, while the whiskers extend to the minimum or maximum of the group or to 1.5x the interquartile range with data points falling outside of that represented as individual points. Symbiont species only represented by a single genome in this dataset are not shown in this figure, as an intraspecies pairwise dissimilarity distance could not be calculated from a single genome.

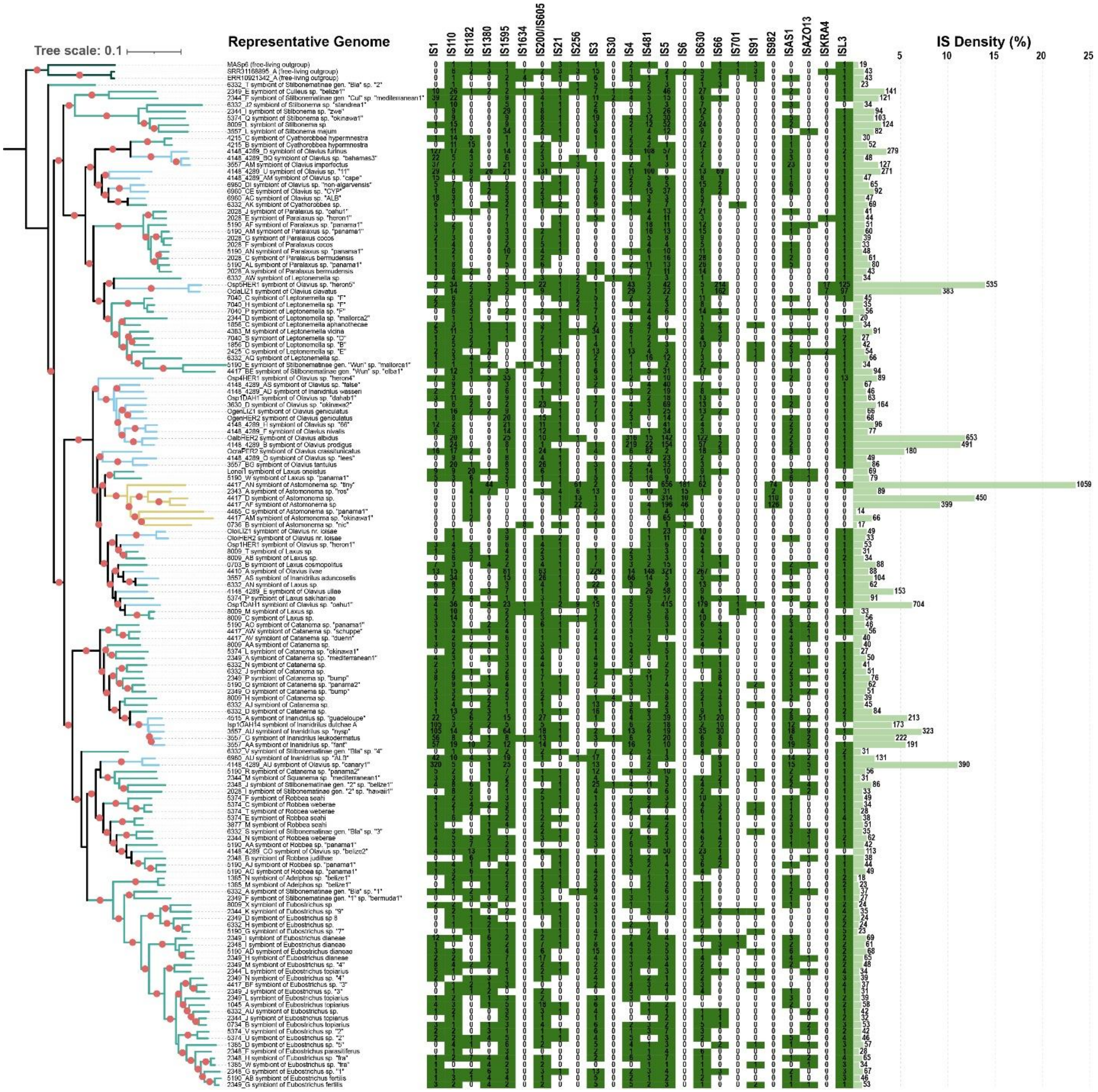


**Figure S3. Phylogenomic tree of *Thiosymbion* and IS family abundance per species.** Phylogenomic tree constructed from single representative genomes for each species. Branch supports are indicated by a red dot (>95%; 1000 bootstraps). IS family presence-absence in representative genome is indicated by the heatmap, with each column representing an IS family and each row corresponding to the adjacent tree leaf. IS family abundance in the representative genome for each species is indicated by the numbers in the heat map. Average IS density (%) per species is indicated by the horizontal green bars and numbers on the bars are the average number of IS sequences per species.

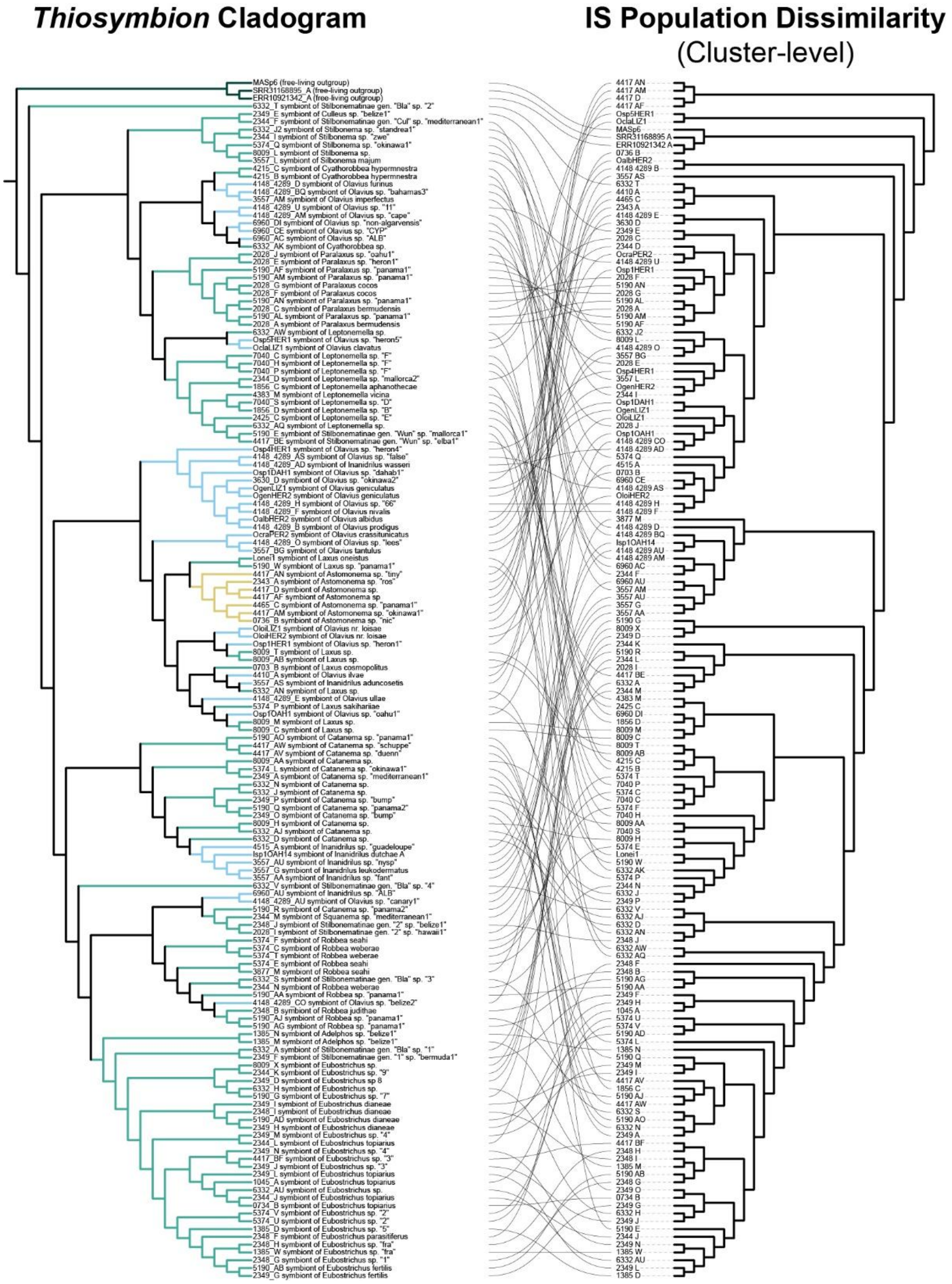


**Figure S4.** Tanglegram of *Thiosymbion* cladogram and IS population subgroup-level dissimilarity (Morisita-Horn) distance matrix represented by UPGMA hierarchical clustering dendrogram.

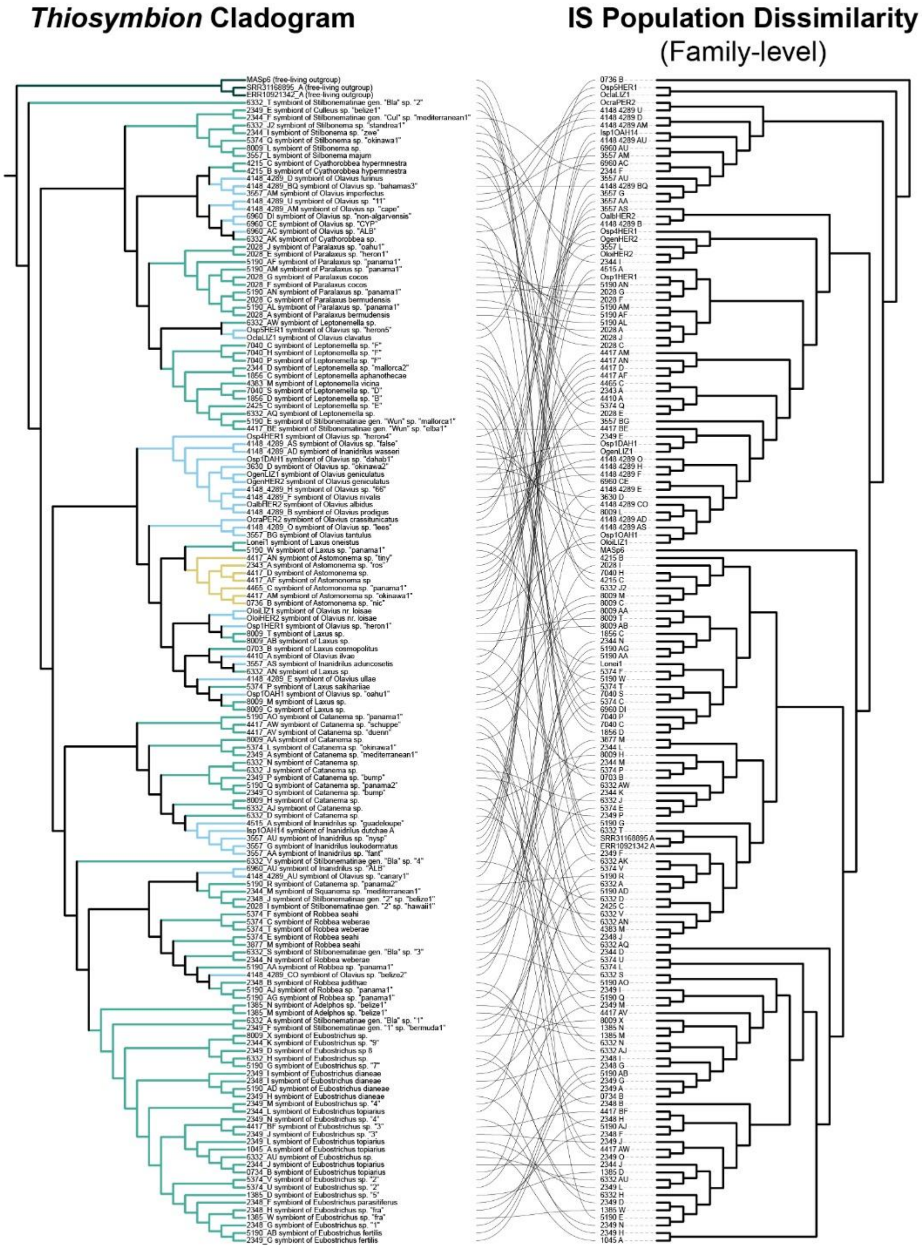


**Figure S5.** Tanglegram of *Thiosymbion* cladogram and IS population family-level dissimilarity (Morisita-Horn) distance matrix represented by represented by UPGMA hierarchical clustering dendrogram.
